# Bio-instantiated recurrent neural networks

**DOI:** 10.1101/2021.01.22.427744

**Authors:** Alexandros Goulas, Fabrizio Damicelli, Claus C Hilgetag

## Abstract

Biological neuronal networks (BNNs) are a source of inspiration and analogy making for researchers that focus on artificial neuronal networks (ANNs). Moreover, neuroscientists increasingly use ANNs as a model for the brain. Despite certain similarities between these two types of networks, important differences can be discerned. First, biological neural networks are sculpted by evolution and the constraints that it entails, whereas artificial neural networks are engineered to solve particular tasks. Second, the network topology of these systems, apart from some analogies that can be drawn, exhibits pronounced differences. Here, we examine strategies to construct recurrent neural networks (RNNs) that instantiate the network topology of brains of different species. We refer to such RNNs as bio-instantiated. We investigate the performance of bio-instantiated RNNs in terms of: i) the prediction performance itself, that is, the capacity of the network to minimize the desired function at hand in test data, and ii) speed of training, that is, how fast during training the network reaches its optimal performance. We examine bio-instantiated RNNs in working memory tasks where task-relevant information must be tracked as a sequence of events unfolds in time. We highlight the strategies that can be used to construct RNNs with the network topology found in BNNs, without sacrificing performance. Despite that we observe no enhancement of performance when compared to randomly wired RNNs, our approach demonstrates how empirical neural network data can be used for constructing RNNs, thus, facilitating further experimentation with biologically realistic network topologies, in contexts where such aspect is desired.

## 1. Introduction

Recent breakthroughs in artificial neural network research have generated a renewed interest in the intersection of biological and artificial neural systems (Richards et al., 2019; Lillicrap et al., 2020; Saxe et al., 2021). This intersection primarily focuses on the biological plausibility of learning rules (Bartunov et al., 2018; Lillicrap et al., 2020) or the similarity of visual stimuli representations in ANNs and BNNs (Cadieu et al., 2014; Güçlü & van Gerven, 2015; Kietzmann et al., 2019). Explicit comparisons of the network topology of BNNs and ANNs are less prominent. This gap primarily exists due to the fact that, despite certain analogies and inspiration derived from neuroscientific findings, groundbreaking advancements in ANNs are driven by engineering goals and computing power (He et al., 2015; Srivastava et al., 2015; Hochreiter & Schmidhuber, 1997) and not by the need for increased neurobiological realism, that necessitates the incorporation of empirical data pertaining to neural network topologies found in nature. Recent studies that examine network topologies of ANNs only focus on generic similarities with biological systems. For instance, building feedforward networks not in a hand-crafted manner, but based on abstract topology models that bear certain similarities with BNNs, such as the Watts-Strogatz model (Watts & Strogatz, 1998) or the Barabási–Albert model (Albert & Barabási, 2002), leads to competitive performances in image classification tasks (Xie et al., 2019). In the context of image recognition, the enrichment of convolutional networks with recurrent connections, mimicking a key feature of BNNs (Markov et al., 2012), leads to competitive performance in benchmark tasks, even outperforming state-of-the-art networks without recurrence (Ming Liang & Xiaolin Hu, 2015). In addition, studies focusing on RNNs, specifically, echo state networks, and evolutionary algorithms applied to ANNs, demonstrate that networks with a modular network architecture, reminiscent of the modular nature of BNNs, exhibit better memory capacity (Rodriguez et al., 2019) and adaptation to new tasks (Clune et al., 2013).

However, such studies have only focused on network topologies that bear generic similarities to BNNs and do not instantiate artificial networks with the actual, empirical network topology that experimental work has unravelled. Despite that, from an engineering standpoint, such level of abstraction may be desired and optimal, it is unknown how empirically-discerned network topology can be incorporated in RNNs and if and to what extent such network topology can lead to beneficial aspects, such as faster learning (fewer training epochs) and better performance (minimization of loss functions in test sets). Examining the effects of empirically discerned network topology om RNNs is important, since ANNs, including RNNs, are increasingly used as models of the brain, and, moreover, the highly structured network topology of animal brains is suggested to serve as a structural prior that is important for rapid learning (Zador, 2019). Notably, despite universal cross-species brain network topology principles, divergence of network topology is also discernible, for instance, when comparing human and monkey brains (Goulas et al., 2014). Therefore, the effects of the incorporation of brain network topology from different species into ANNs should be examined. Here, we explicitly address this gap. Instead of examining network topologies that are biologically inspired or analogous to BNNs, we build RNNs that are bio-instantiated, that is, they embody the empirically discerned network topology found in natural neural systems, that is, monkey and human brains. This is a necessary step to explicitly examine the intersection between ANNs and BNNs at the network topology level. We examine the impact of bio-instantiated RNNs in a series of working memory tasks that entail the ability to track relevant information as a sequence of events unfolds across time. Working memory is a key cognitive capacity of biological agents, extensively studied in cognitive psychology and neuroscience (Conway et al., 2003). Moreover, such capacity is also desired in engineering tasks where sequential data are processed.

Since RNNs are hand-engineered and BNNs are a product of evolution, apart from intuitive expectations, it is not clear how network topology of BNNs can be incorporated into RNNs and how such network topology impacts the performance of RNNs. Our main goal is to investigate different *strategies to construct bio-instantiated RNNs from empirical data on BNNs* and elucidate the *performance of these bio-instantiated RNNs*, for instance, assess if BNN-based network topology is a potentially advantageous structural prior that would situate the system at hand in an advantageous starting point, that is, effecting performance and speed of training when the networks is subsequently trained.

Our contributions are as follows: we investigate three different strategies to construct bio-instantiated RNNs from experimental observations of the human and monkey brain networks and the impact of such strategies on the performance and speed of learning of the bio-instantiated RNNs. We examine these aspects in the context of a commonly used learning algorithm, that is, backpropagation-through-time, and rate models with continuous activation functions. Our results indicate that not all strategies for creating bio-instantiated RNNs lead to the same performance. A lack of enhancement of the performance of the bio-instantiated RNNs when compared to randomly wired RNNs showcases the need for further modifications, possibly encompassing more biologically realistic learning algorithms and activation functions, conjointly with the incorporation of additional dimensions across which the performance of the bio-instantiated RNNs should be evaluated.

It is important to highlight the distinction between *network architecture* and *network topology*, as currently used in this report. State-of-the-art RNNs, such as LSTMs (Hochreiter & Schmidhuber, 1997), exhibit a non-random engineered architecture (different types of interconnected gates), but a random topology (all-to-all connectivity with randomly initialized weights). In addition, the classic Elman networks (Elman, 1990) exhibit a non-random architecture, for instance, the input layer connects to the hidden recurrent layer, but not directly to the output layer. However, the topology of the hidden recurrent layer is all-to-all, and, thus, in stark contrast to the network topology of biological systems, such as the human and monkey brain network.

### 1.1. Related work

In the realm of neuroscience, experimental work and network analysis revealed that BNNs, such as the worm neural network, have a characteristic network topology, that is, neurons in BNNs do not connect in a random or all-to-all fashion, but exhibit preferential connectivity that obeys certain wiring principles (Varshney et al., 2011; Motta et al., 2019). Comprehensive, single neuron connectivity at a large-scale, whole-brain level in animals, such as humans, is currently experimentally intractable. While such information is still lacking, detailed experimental data reveal how neuronal populations in different brain regions of human and non-human animals are wired to each other. These experimental observations highlight that neuronal populations inhabiting the various regions of animal brains, do not connect to each other in a random or all-to-all fashion, but exhibit preferential connectivity and connection weights, thus, forming a characteristic network topology (van den Heuvel et al., 2016; Goulas et al., 2019). In sum, the network topology pertaining to a plethora of BNNs is in contrast to the hand-crafted engineering-driven network topology of ANNs. A recent study examines the effect of constructing RNNs with the empirically discerned topology of the human brain network (Suarez et al., 2020). This study, however, examines a different class of RNNs (echo state networks) than the one that we examine in our approach (Elman networks), and in addition, echo state networks are trained with a different algorithm as the one that is used here (backpropagation-through-time). Moreover, our approach is comparative, that is, it uses empirical brain network data from different species, thus, conveying additional conceptual and methodological advantages (see Discussion).

In the realm of ANN research, the characteristic, non-random network topology of BNNs has started to gain attention. In the context of evolutionary optimization, networks that are optimized for minimization of their connection cost, that is, the physical distances spanned by connections between the neurons of the system, as observed in BNNs (van den Heuvel et al., 2016; Goulas et al., 2019), also exhibit modularity, a network topology feature that also pertains to BNNs (Clune et al., 2013). Moreover, networks that are evolved with the connection cost incorporated in the objective function, are characterized by enhanced performance and adaptation capacity to new tasks (Clune et al., 2013). However, such approach *evolves a network architecture* based on biologically grounded principles and does not examine the *impact of the network topology of BNNs* if directly embedded in RNNs that are subsequently trained with a *widely used learning algorithm for RNNs*, that is, backpropagation-through-time. Moreover, incorporating a key general feature of BNNs, that is, the fact that connections between neurons are reciprocal, led to convolutional neural networks with recurrence that outperform purely feed-forward convolutional networks in benchmark object recognition tasks (Ming Liang & Xiaolin Hu, 2015). Such hybrid convolutional networks with recurrence have been further enriched and tested in a neuroscientific context, leading to better fit with empirical data recorded from the human brain during object recognition (Kietzmann et al., 2019). While such studies showcase the potential benefits of crafting ANNs with network topology features found in BNNs, nevertheless, they embody a general wiring feature, in this case recurrence, and do not embody the exact recurrent patterns of connections between brain areas in the way that empirical data indicate. In addition, (You et al., 2020) introduce a BNNs to ANNs conversion strategy that relies on the concept of relational graphs, but such approach uses network topology of BNNs as a generator for feedforward networks and not for the creation of RNNs with a biologically-based network topology. Lastly, at the interface of neuroscience and ANNs, RNNs have been used to explore diverse tasks, including higher-order cognitive domains (Eliasmith et al., 2012; Yang et al., 2019; Cueva et al., 2020), but such experiments *do not use RNNs that have a neurobiologically realistic network topology*.

## 2. Constructing bio-instantiated RNNs

All the steps and strategies for extrapolating from empirical data on BNNs to RNNs are possible with the *bio2art* package: https://github.com/AlGoulas/bio2art

### 2.1. Faithful extrapolations from empirical data

Analysis of empirical data demonstrates that BNNs are characterized by key network topology principles dictating how the neuronal populations of the different brain regions connect to each other. Such network topology principles include preferential non-random connections between specific brain regions and non-random connection weights (Bullmore & Sporns, 2009; van den Heuvel et al., 2016; Goulas et al., 2019) (Fig. 1 A). Here, we exploit this plethora of empirical evidence to construct the network topology of the hidden recurrent layer of an RNN, specifically an Elman network (Elman, 1990). We should note that our study focuses on the ramifications of the incorporation of network topology found in nature into RNNs, and not solving engineering problems *per se*. Thus, despite the obvious and known limitations of Elman-type RNNs (Hochreiter & Schmidhuber, 1997), we chose this type of RNNs due to their suitability to test the key question at hand, that is, the effect of biological versus random network topology in artificial systems. Note that the present framework of converting BNNs to RNNs is general enough to accommodate other types of RNNs, such as LSTMs (Hochreiter & Schmidhuber, 1997) and GRU networks (Cho et al., 2014).

**Figure 1.**
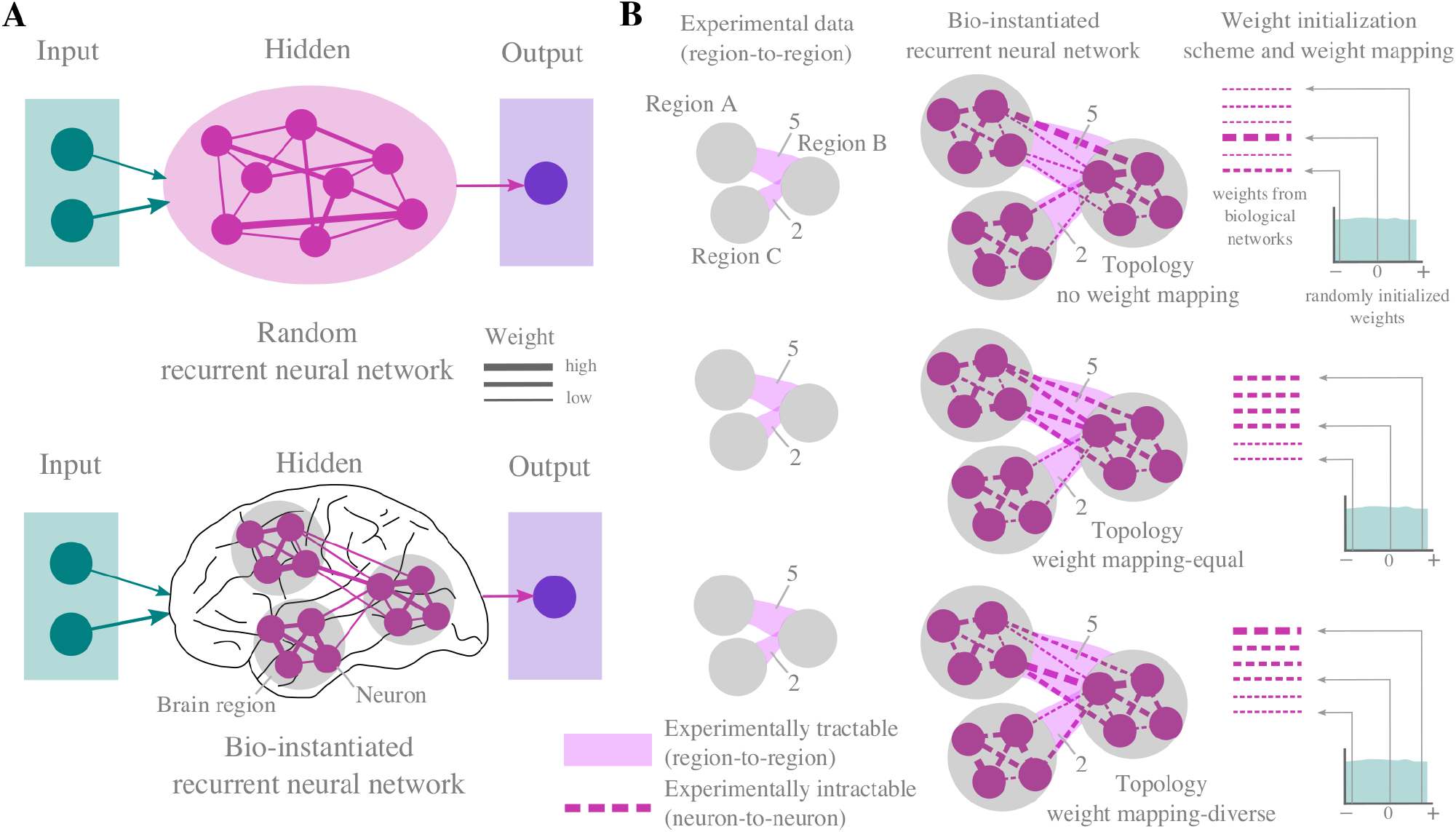
Constructing bio-instantiated RNNs. A. RNNs have a random or all-to-all topology, with neurons connecting to each other with a random weight sampled from e.g., a uniform distribution. On the contrary, BNNs do not exhibit a random or all-to-all topology. Instead, neuronal populations within a brain region connect to specific neuronal populations in other brain regions. In addition, brain regions, and the underlying neurons within this region, do not connect to each other with random weights. Instead, region-to-region, and, presumably, the underlying neuron-to-neuron connections, have non-random connection weights. B. Extrapolating from empirical data on biological neural networks to construct bio-instantiated RNNs. Empirical data on brain networks offer quantitative data on region-to-region connection weights (pink solid lines). Neuron-to-neuron connection weights are computed in such a way that the sum of all neuron-to-neuron connection weights between two regions (e.g., region A and B) is equal to the empirical region-to-region connection weight (region A to region B=5). In total three strategies were followed to construct the bio-instantiated RNNs from empirical data: *topology-no weight mapping, topology, weight mapping-equal, topology weight mapping-diverse*. See sections 2.1–2.3 for details. Note that this schematic drawing depicts the human brain, but the same strategies and principles apply to the monkey brains.

#### 2.1.1. From regions to neurons

One first challenge in translating empirical BNNs to RNNs is the fact that the wiring of biological system, such as the human and monkey brain, is not experimentally tractable at the neuron-to-neuron level, but summarized as a brain region-to-brain region wiring diagram (Fig. 1 B). For brains such as the monkey brain, the wiring diagram is based on a fixed number of regions derived from a brain parcellation that currently encompasses 30-60 brain regions (Markov et al., 2012; Majka et al., 2020). Thus, *a first challenge is to devise a way to extrapolate from this fixed number of network size and create RNNs that obey the empirical brain region-to-brain region network topology, but can have an arbitrary size*. An arbitrary size, that is, a RNN with an arbitrary number of neurons in the hidden layer, is desired, since different tasks have different demands in terms of the network size that is adequate for achieving competitive performance. Currently we offer a simple solution: for each brain region, we assume that it is populated by N neurons. For instance, in the example illustrated in Fig. 1, the empirically discerned network used as an example is composed of three regions (region A, B and C) and each region contains N=4 neurons. Note that for the current experiments, bio-instantiated RNNs were also constructed by assuming that each region contains 4 neurons.

#### 2.1.2. Region-to-region and neuron-to-neuron wiring

A second challenge is to translate the region-to-region wiring diagram to a neuron-to-neuron wiring diagram. The empirical evidence dictate that at the *binary topological level*, region A and B are connected, as well as region B and C, but not region A and C (Fig. 1 B). We populate each region with four neurons. The neurons within each region connect to the neurons of another region based on the empirical data, for instance, neurons within region A connect to neurons in region B, but not region C.

Moreover, empirical data on brain networks indicate that the strength of connections is heterogeneous. In the example in Fig. 1 B), the weight of the connection between region A and B is 5, whereas the weight between regions B and C is 2. Thus, from a *weighted topological level* standpoint, we extrapolate connection weights between the neural populations of the different regions. We should note that in the current setup, the number of connections that a neuron can form is controlled by a parameter dictating the percentage of connections that a neuron will form, out of the total number of connections that can be formed. Here, we set this parameter to 0.8, that is, 80% of all possible connections are formed. In the example depicted in Fig. 1 B, with this parameter set to 0.8, each neuron in region A connects to 3 neurons in region B (note, however, that only two connections are depicted for simplifying the visualization).

#### 2.1.3. Within region neuron-to-neuron wiring

Empirical, quantitative and comprehensive data for neuron-to-neuron connectivity within each brain region are lacking. However, existing empirical data suggest that within-region connectivity strength constitutes approximately 80% of the extrinsic between-region connectivity strength (Markov et al., 2012). Therefore, the intrinsic, within-region connectivity in our BNN to RNN translation strategy followed this rule. Thus, with this rule, in the example depicted in Fig. 1 B, the total strength of connections between neurons in region A is 80% of 5, that is, 4. It should be noted that the number of connections that a neuron can form with neurons of the same region is controlled by a parameter dictating the percentage of connections that a neuron forms, out of the total number of connections that can be formed. Here we set this parameter to 1, that is, all connections between neurons within a region are formed. Thus, in the examples in Fig. 1 B, within a region, e.g., region A, all neurons connect to all neurons within the same region.

#### 2.1.4. Assigning region-to-region connection weights to neuron-to-neuron weights

As mentioned, current empirical data at the whole brain level offer quantitative information only for region-to-region connection strength. We use this empirically discerned information to assign neuron-to-neuron connection weights. This necessarily entails extrapolations. Two ways of assigning neuron-to-neuron connection weights from region-to-region weights were adopted. First, given connection weight M between two regions, and K neuron-to-neuron connections between the two regions, all neuron-to-neuron connections have an equal weight calculated as M/K. We refer to this scheme as the *equal* scheme. In the example in Fig. 1 B (*middle panel:weight-mapping equal*), neuron-to-neuron connections between region B and C have a weight equal to 1, since the region-to-region weight is 2 and we have in total 2 neuron-to-neuron connections. Second, given connection weight M between two regions, and K neuron-to-neuron connections between the two regions, the i^th^ neuron-to-neuron connection weight is estimated as w_i_, where w_i_ is a subset of W, and i=1,2,…K, with sum(W)=M. In other words, M is expressed as a sum of K partitions, where K=number of neuron-to-neuron connections between two regions. We refer to this scheme as the *diverse* scheme. In the example in Fig. 1 B (*bottom panel:weight-mapping diverse*), neuron-to-neuron connections between region B and C have e.g, weights equal to 1.5 and 0.5, since the region-to-region weight is 2 and we have in total 2 neuron-to-neuron connections. Note that extrapolating from region-to-region to neuron-to-neuron weights involves extrapolations and, in the *diverse* scheme described above, a degree of randomization is entailed: while the sum of connection weights between neuronal populations between two regions is constrained and dictated by empirical network data, the individual neuron-to-neuron weights are random partitions of the empirical weight as described above.

### 2.2. Mapping neuron-to-neuron weights and RNN weights

Weights of RNN involve an initialization so that the RNN can be trained properly. Here, we used a default initialization scheme where weights were initialized from a uniform distribution 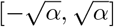 where *α*=1/NN, and NN the total number of neurons of the hidden layer of the RNN. We should note that other suggested initialization schemes for RNNs, for instance, as described in (Le et al., 2015), were not applicable in our case, since they require that the recurrent hidden layer is the identity matrix and the very purpose of our study is to examine the impact of the incorporation of the network topology dictated by the network topology of BNNs into RNNs and this topology is clearly not an identity matrix. Alternative weight initialization schemes that are usually adopted, that is He (He et al., 2015) and Xavier (Glorot & Bengio, 2010) initialization schemes, led to qualitatively the same results. We should note that instead of using a uniform distribution, possible extensions may involve weights that can also be initialized by sampling from families of distributions based on certain empirical observations on neuron-to-neuron connection weights (Motta et al., 2019).

An important step is to assign the weights that were initialized from a uniform distribution 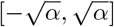 to the neuron-to-neuron weights that were extrapolated from empirical neural network data (Sections 2.1.1-2.1.4). Here, we adopt a simple approach. We rank-order the neuron-to-neuron weights that were extrapolated from empirical neural network data and the weights that were initialized from a uniform distribution. Subsequently, each neuron-to-neuron weight is assigned to the corresponding rank-ordered weights initialized from the uniform distribution. In this way, the higher neuron-to-neuron weights extrapolated from the empirical data are assigned to the higher weight values that were initialized from a uniform distribution Fig. 1 B (*middle panel:weight-mapping equal and bottom panel:weight-mapping diverse*). Note that this matching of the rank of weights that were initialized from a uniform distribution to the neuron-to-neuron weights that were extrapolated from empirical neural network data does not apply to the *topology no weight mapping* strategy (Fig. 1 B *top panel*), since in this case the topology of the bio-instantiated RNN is only constrained from biological data at the binary level.

We should note that with this weight mapping scheme, the weakest neuron-to-neuron weights extrapolated from empirical data are assigned to the lower values of the weights that were initialized from a uniform distribution, weights that include negative values. This is an assumption with no rigid empirical evidence and could be accommodated differently in the future (see Discussion). This weight mapping leads to bio-instantiated RNNs that are suitable to be trained with commonly used training algorithms, that is, backpropagation-through-time.

### 2.3. Bio-instantiated RNNs

The steps described in sections 2.1 and 2.2 allow building RNNs that exhibit a network topology dictated by the network topology of BNNs, both at the binary topological level (what neuron connects to what neuron) and weighted topological level (how strongly two neurons connect to each other). Here, we focus on a simple RNN, that is, an Elman network (Elman, 1990), defined as:

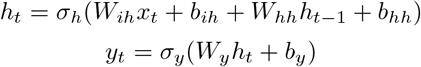

with *x_t_*, the input vector, *h_t_* the hidden layer vector, *y_t_* the output vector, *W_ih_*, *W_hh_* and *W_y_* the weight matrix for the input, hidden and output layers, respectively and *σ_h_*,*σ_y_* the respective activation functions and *b_ih_*, *b_hh_*, *b_y_* the respective bias terms.

The bio-instantiated RNNs are, thus, Elman networks, and their bio-instantiated nature is based on the instantiation of the weight matrix *W_hh_* from empirical data on the topology of BNNs as described in sections 2.1 and 2.2. Based on these descriptions, we construct the bio-instantiated RNNs based on different strategies (summarized in Fig. 1 B). Specifically, we adopt three strategies: *topology no weight mapping, topology weight mapping-equal, topology weight mapping-diverse*, and the weight matrix *W_hh_* in these strategies is defined as follows. For the *topology-no weight mapping* strategy: *W_hh_* = *W_bio_* ⵙ *W_init_*, with *W_bio_* the recurrent matrix extrapolated from the empirical network data with the *topology no weight mapping* strategy, *W_init_* a recurrent matrix with randomly initialized weights, as described in Section 2.2, and ⵙ the Hadamard product. Thus, *W_hh_* is a recurrent matrix with randomly initialized weights that exhibits the non-random binary topology dictated by *W_bio_*, but no relation between weights w ∈ *W_init_* and weights *w′ ∈ W_bio_*. For the *topology weight mapping-equal* and *topology weight mapping-diverse* strategy, *W_hh_* = *W_bio_* ⊙ *W_init_* with weights w ∈ *W_init_* and weights *w′ ∈ W_bio_*, and *ρ*(*w, w′*) = 1, with *ρ* denoting Spearman’s rank correlation. Thus, in these strategies, both the *binary and weighted* topology of the bio-instantiated RNNs is shaped by the topology of the BNNs. Note also that in these two strategies, the weights of *W_bio_* and *W_init_* are mapped in the same way just as described above, but the difference between these two strategies is the extrapolation of the weights from empirical observations on weights of BNNs (see section 2.1.4).

Note that the weights of the bio-instantiated RNNs are subject to modifications, that is, weight updates during the training process (see section 4). During this process, only the non-zero weights are modified, that is, exisitng connections. It is suggested that the network topology of animal brains is highly structured and such feature may result in faster learning (Zador, 2019). Thus, in our approach the aim of the different strategies to create bio-instantiated RNNs is to examine if the empirical network topology configurations can convey an advantageous starting point for the networks to be trained compared to networks that do not embody biological network topology features. Note that the input connections from the input to the hidden layer, as well as the output connections from hidden to the output layer, are all-to-all, that is, there is not preferential input to, for instance, the neurons of “vision areas” of the hidden layer, or preferential output connectivity from neurons in the “motor-areas” of the hidden layer to the output layer. Since each neuron belongs to a brain area that corresponds to the actual biological system (monkey or human brain), the aforementioned specificity of input and output connections can be incorporated in the future, due to the flexibility of our framework. Note as well that the results reported here are converging with results that use bio-instantiated Echo State Networks (Damicelli et al., 2021).

## 3. Tasks

Our approach is motivated by neurobiology, and, thus, we focus on a cognitive domain that is commonly used in experimental neuroscience. Hence, we tested the working memory capacity of the RNNs. The choice of this capacity is motivated by a key property of RNNs and biological systems, that is, the capacity to remember information that is not directly present at a given time point, but relevant for the execution of a task. The importance of working memory is highlighted, for instance, in humans, where working memory capacity is related to general intelligence (Conway et al., 2003) and many tasks with an engineering nature and data with a sequential nature. Thus, we examine the performance of the networks in the context of two working memory tasks: sequence memory (seq mem) and nback memory (nback mem) Fig. 2. Naturally, the repertoire of tasks can be extended in the future to other domains and also encompass benchmark tasks that are used for engineering-oriented approaches.

**Figure 2.**
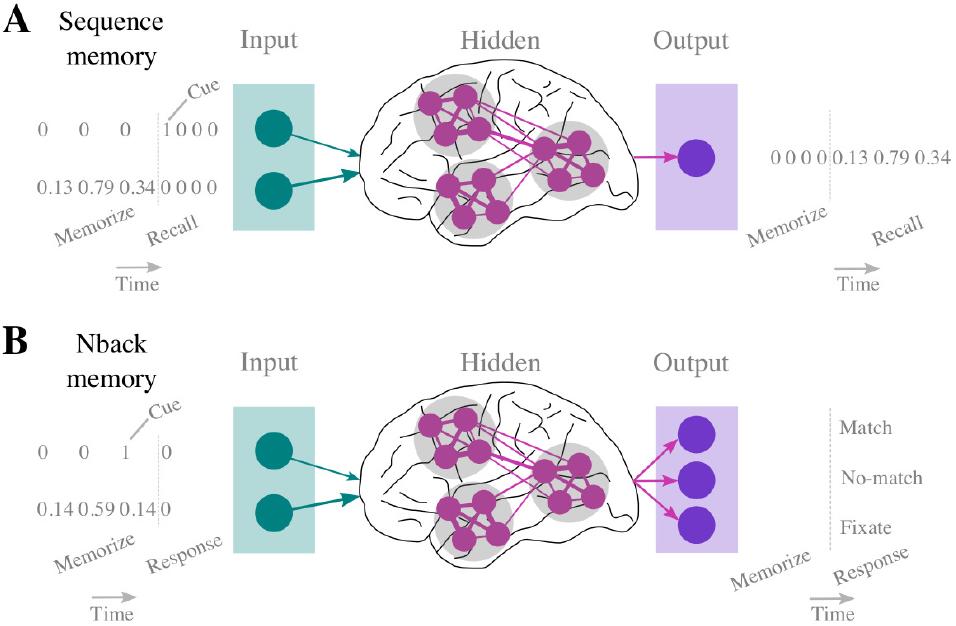
Working memory tasks. A. Sequence memory task. In this task, a sequence of N numbers has to be memorized and recalled (see section 3.1 for details). B. Nback memory task. In this task, a number has to be matched to a number n time steps ago (see section 3.2 for details).

### 3.1. Sequence memory

For the sequence memory task, a sequence of N numbers was generated uniformly and randomly from the [0 1) interval and fed into the artificial neural network (memorize phase). When a cue signal was provided, denoted by “1” fed into a separate input neuron from the one used as input for the sequence of N numbers, the network had to generate the exact sequence of N numbers (recall phase) Fig. 2 A. The mean square error (MSE) between the predicted and actual sequence of numbers served as the loss function for this task. It should be noted, that since we were interested in the memory capacity of the network, the loss was computed only on the output of the network during the recall phase.

### 3.2. Nback memory

For the nback memory task, a sequence of M numbers was generated uniformly and randomly from the [0 1) interval and fed into the artificial neural network (memorize phase). When a cue signal was provided, denoted by “1” fed into a separate input neuron from the one used as input for the sequence of M numbers, the network had to respond by judging if the last number in the sequence is the same as the number n time steps ago (response phase) Fig. 2 B. The responses of the network define three classes: fixate (no response), match, or no-match Fig. 2 B. The negative log-likelihood between the correct class/response and the network output served as the loss function for this task. It should be noted, that since we were interested in the memory capacity of the network, the loss was computed only on the output during the response phase.

## 4. Experiments

The code for running the experiments with the aforementioned RNNs and tasks is available at https://github.com/AlGoulas/bio_rnn_mem

### 4.1. Training and experimental parameters and performance measures

We created bio-instantiated RNNs with three strategies: topology no weight mapping, topology weight mapping-equal and topology weight mapping-diverse. This procedure was applied to three different species: marmoset monkeys, macaque monkeys and humans. The networks were trained with backpropagation-through-time. We should note that the goal of the experiments was not to select the optimal combination of hyperparameters, but to elucidate the impact that the different strategies of creating bio-instantiated RNNs have on the prediction capacity and speed of training of the networks. Therefore, in order to observe the impact of the biological topology across different settings, experiments were carried out with a range of varied and fixed parameters. The learning rate was set to 1 × 10^*−*4^. We ran the experiments with two different commonly used optimizers, that is, Adam and RMSprop. We ran the experiments with two activation functions, that is, ReLU and tanh. Lastly, we varied the parameter of the memory task to be executed, that is, for the sequence memory task, the length N of the sequence of numbers to be memorized was varied (N=3,5,10) and for the nback memory task, the number of time steps n that were needed to compare the current target number to was also varied (n=2,3,4). Thus, in total, 12 unique combinations were generated from the aforementioned parameter space for each species and each strategy for creating bio-instantiated RNNs. For each of these cases, the networks were trained in 5 separate runs. For computational reasons, the epochs for training were set to 500 for the sequence memory task and 3000 for the nback task. For each task, 1000 trials were generated and 80% of these trials were used for training the network and the rest were used for estimating the test loss that was used as a performance measure of the network. Specifically, the goal of the experiments was to examine the impact that the network topology found in biological systems has on the prediction capacity and speed of training of the RNNs. Therefore, the loss on test trials was used as a measure of prediction capacity and the epoch where the minimum loss was observed was used as a measure of how fast the network reached its maximum performance.

### 4.2. Random network topology as control cases

Since the topology of RNNs is the focus, as control cases, the experiments, as described above, took place, but now with bio-instantiated RNNs with a topology that was subsequently randomized. This randomization of the topology entailed shuffling the connection weights of the bio-instantiated network, generated from each strategy described in Fig. 1, thus, destroying any structured topology imposed by BNNs. Note that results obtained from these randomized RNNs are prefixed with “`r_`”.

## 5. Results

### 5.1. Effects of topology on loss minimization

For the sequence memory task, all strategies for constructing bio-instantiated RNNs led to a comparable performance, except one strategy, that is, topology weight mapping-equal (`b_w_eq` in Fig. 3), that led to higher median loss or more excessive variance of minimum loss, thus, less consistent convergence to the minimum loss across parameters and experimental runs. What is of note is that all the other strategies of constructing bio-instantiated artificial networks led to a comparable performance to artificial networks with random topology (as is commonly the case with RNNs) (Fig. 3). This held true for topologies constructed from brain network data of monkeys and humans.

**Figure 3.**
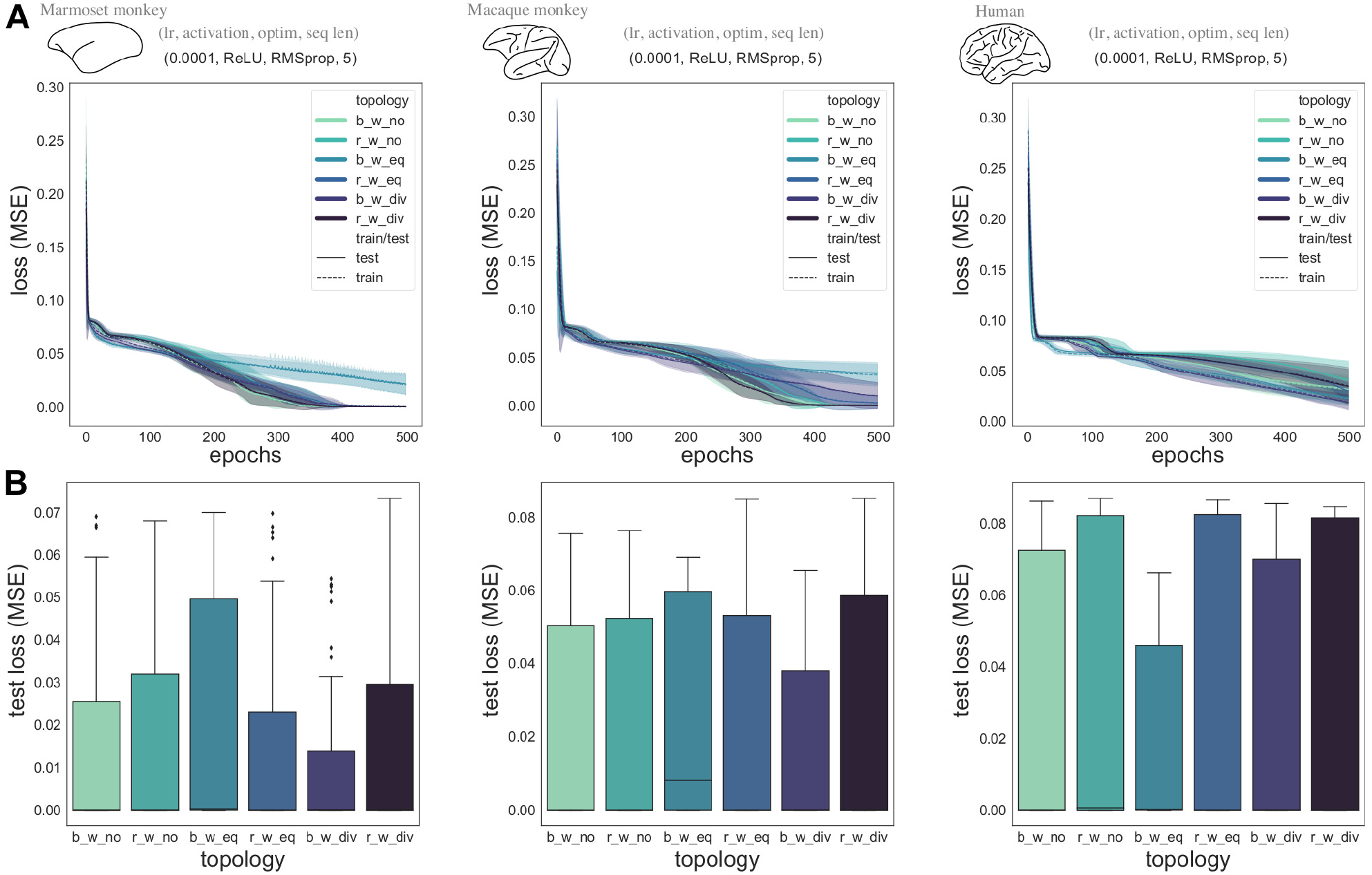
Sequence memory task results. A. Evolution of the loss function (MSE) across epochs for the different strategies, resulting in different topologies, across different species. Results are depicted for one parameter combination, denoted by the tuple (lr, activation, optim, seq len), denoting the learning rate, activation function, optimizer and sequence length for the sequence task that was used. Shaded area denotes the standard deviation across the different runs (=5) of the experiments. B. Summary of the loss on test data across all parameter combinations for the different topologies and species. Abbreviations: `b_w_no`=biological topology - no weight mapping, `r_w_no`=random topology - no weight mapping, `b_w_eq`=biological topology - weight mapping-equal, `r_w_eq`=random topology - weight mapping-equal, `b_w_div`=biological topology - weight mapping-diverse, `r_w_div`=random topology - weight mapping-diverse. These abbreviations correspond to the conversion strategies described in Fig. 1.

For the nback memory task, the topology weight mapping-equal (`b_w_eq`) strategy led to the worse observed performance across different parameters and across networks constructed from different brain networks (Fig. 4). For all the other strategies that were followed to construct bio-instantiated RNNs, a comparable performance, that is minimum test loss, was observed when compared to RNNs with random topology (Fig. 4). Therefore, these results showcase which strategies can be followed to construct RNNs with neurobiological network topology, without sacrificing performance. This held true for all the brain networks, that is, monkey and human brain networks, used for creating the artificial counterparts.

**Figure 4.**
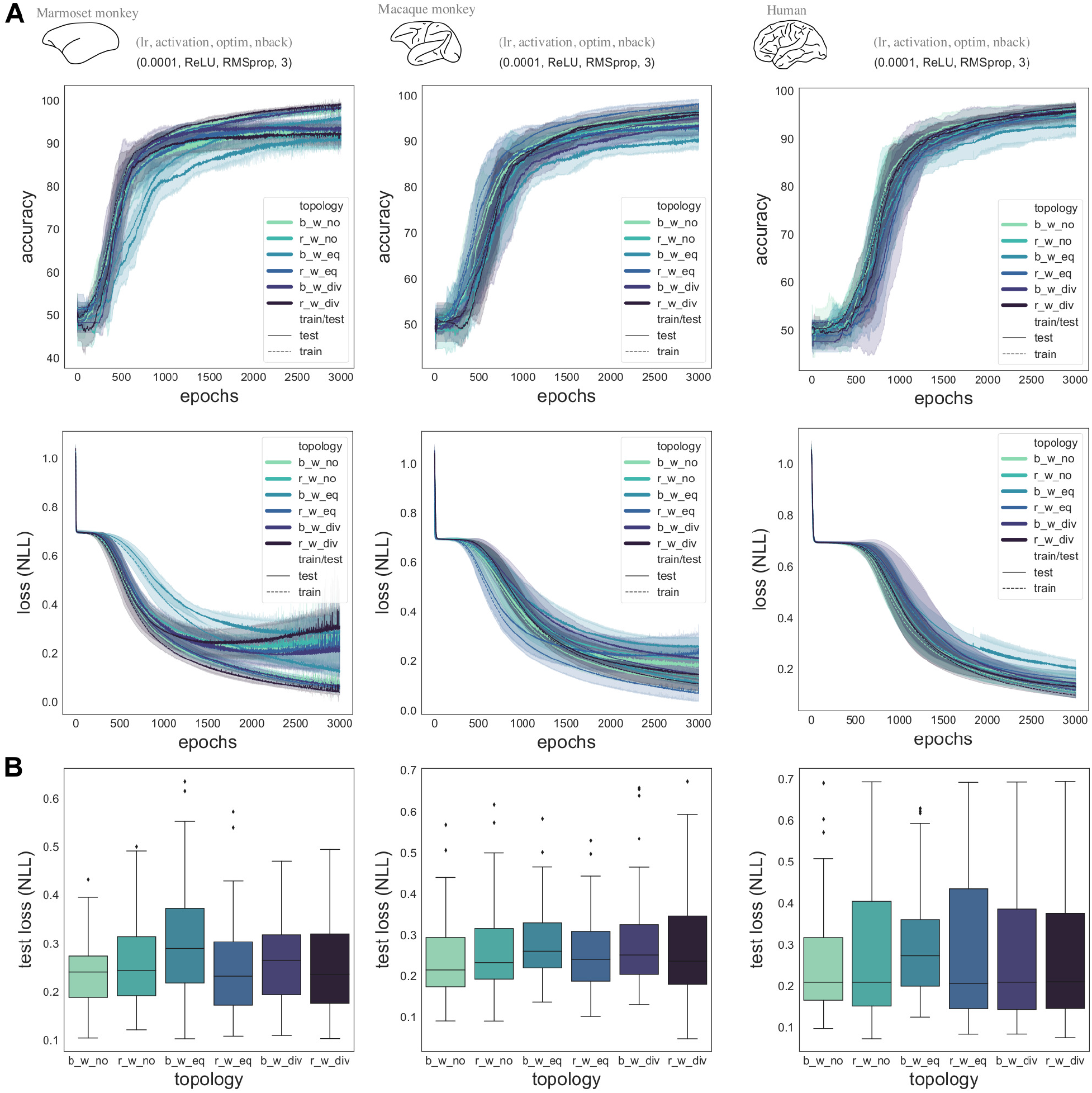
Nback memory task results. A. Evolution of the loss function (NLL) and the accuracy across epochs for the different strategies, resulting in different topologies, across different species. Results are depicted for one parameter combination, denoted by the tuple (lr, activation, optim, seq len), denoting the learning rate, activation function, optimizer and sequence length for the sequence tasks that was used. B. Shaded area denotes the standard deviation across the different runs(=5) of the experiments. Summary of the loss on test data across all parameter combinations for the different topologies and species. Abbreviations as in Fig. 3.

It should be noted that for both tasks no difference was observed between the strategy that takes into account only the binary topology of BNNs (what neuron connects to what neuron, `b_w_no`) and the strategy that also takes into account the weights (what neuron connects to what neuron and with what weight, `b_w_div`). This indicates that, in the current context, constructing RNNs based on the binary or weighted network topology of BNNs does not impact performance.

### 5.2. Effects of topology on speed of training

Apart from examining the effect of topology and the different strategies for constructing bio-instantiated RNNs on prediction capacity, we also examined the effect on speed of training, that is, how fast the network can reach its optimal performance (minimum loss). Once again, the topology weight mapping-equal (`b_w_eq`) strategy led to certain cases where slower learning across epochs was observed, that is, the best performance (minimum test loss) was achieved in late epochs compared to what is the case for their randomized controls (r_w_eq) and the rest of the biological to artificial conversion strategies (Fig. 5). Such cases, where sub-optimal speed of training was observed for the topology weight mapping-equal strategy, involved all BNNs, that is, the monkey and human brain networks. Hence, these results dictate what biological to artificial conversion strategies can lead to RNNs with a biological topology without any loss in speed of training.

**Figure 5.**
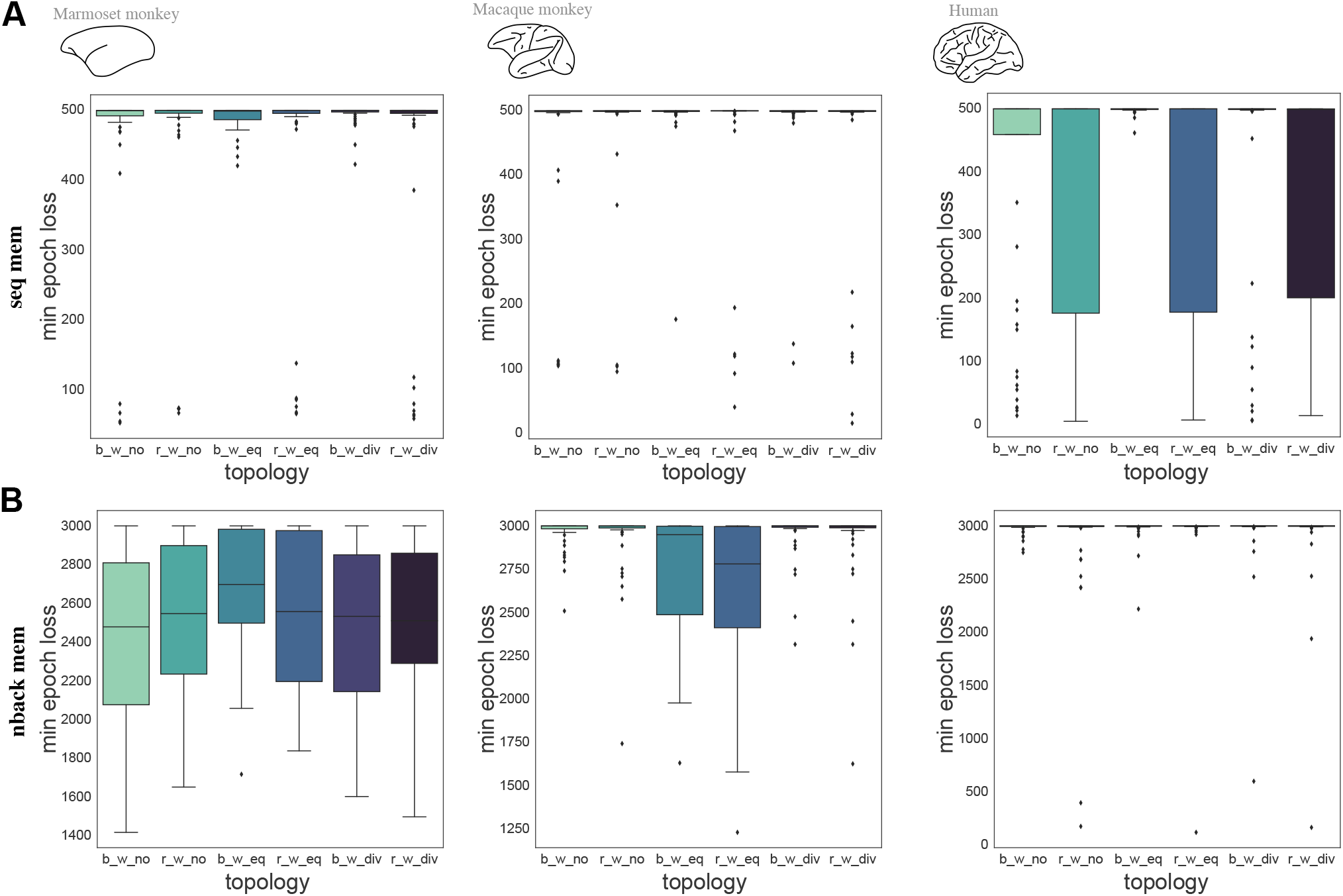
Epoch where the minimum loss on tests data is achieved for A. the sequence memory task and B. the nback memory task, across different species. Abbreviations as in Fig. 3.

As it was the case with prediction capacity, for both tasks, no difference in the speed of training was observed between the strategy that takes into account only the binary topology of BNNs (`b_w_no`) and the strategy that also takes into account the weights (`b_w_div`). This indicates that, in the current context, constructing RNNs based on the binary or weighted network topology of BNNs does not impact speed of training.

## 6. Discussion

### 6.1. Examining the network topology of ANNs and BNNs beyond the lens of abstract analogies

We complement existing approaches emphasizing network topology (Ming Liang & Xiaolin Hu, 2015; Kietzmann et al., 2019; You et al., 2020) by offering a novel perspective, that is, creating bio-instantiated RNNs that are based on empirical data on the topology of multiple biological neural systems, that is, the marmoset and macaque monkey brain network, as well as the human brain network. Such an approach is necessary to explicitly elucidate postulations on commonalities of ANNs and BNNs at the network topology level and offer concrete evidence related to suggestions that innate cognition and fast learning pertaining to animals is based on the particular network topology of brain networks (Zador, 2019). Our results show that a network topology based on BNNs does not serve, *in the current model setup and mode of evaluation*, as an advantageous structural prior positively effecting performance during the subsequent training of the network. However, our results show how we can construct RNNs with a biology-based network topology, without sacrificing performance, a fruitful avenue for approaches that need enhanced neurobiological realism (e.g., RNNs as models for the brain). Specifically, our experiments demonstrate that not all strategies for creating bio-instantiated RNNs from empirical network data lead to the same performance. Specifically, the strategy that performs worse is based on the assumption that connection weights between neuronal populations inhabiting two different brain regions are equal. On the contrary, strategies that assume that such weights are diverse, that is, they have heterogeneous values, are achieving the best observed performance in the currently examined experimental setup. Notably, the sparse empirical evidence on brain connectivity of mammals at a single neuron-to-neuron level indicate that connection weights between neurons are diverse (Motta et al., 2019). While many different functional advantages can be conveyed by such heterogeneous weights of connections, in the current context, these heterogeneous connections may be beneficial to maximize the diversity of non-linear transformations to the input of each neuron and, thus, achieve a more diverse set of outputs per neuron, and, thus, potential inputs to each neuron, since the output of each neuron is multiplied by diverse, and not equal, weights pertaining to its outgoing connections. Such diverse output and input would facilitate the statistical independence between the activity of the neurons in the RNN, a feature that, for instance, in the context of Echo State Networks appears to be crucial for the enhanced performance of the system (Morales et al., 2021). Notably, statistical independence is also considered a key feature of BNNs, that is, the cerebral cortex (Barlow & Földiak, 1989).

From a broader standpoint, our approach, explores the effects of different strategies of converting data of diverse BNNs to RNNs, via a tool that is freely available (Goulas, 2020), thus, facilitating further experimentation. For instance, a novel comparison of the human ventral visual system and deep convolutional neural networks (Güçlü & van Gerven, 2015) is now feasible by situating this comparison on a biologically realistic plane that takes into account the empirical data that describe the exact network topology of the biological system of interest. Importantly, our approach uses data at a whole brain level, and does not only focus on a particular system, thus, enabling an arbitrary set of investigations that encompass any system and cognitive and behavioral domains of interest.

### 6.2. Binary and weighted topology

Innate cognition and fast learning pertaining to animals may be based on the non-random network topology of the brain (Zador, 2019). Brain networks exhibit a non-random topology both at the binary and weighted level (Markov et al., 2012; Goulas et al., 2019). Evolutionary algorithms applied to ANNs that do not allow optimization of weights, but only optimization of topology at the binary level, lead to competitive performance on a classification benchmark (Gaier & Ha, 2019). Thus, topology of ANNs at the binary level also encodes “know-how” for classification tasks. In our case, we investigated the significance of brain network topology of diverse animals as a potentially advantageous structural prior that would render the system at hand more efficient in terms of performance and speed of training when subsequently trained with backpropagation-through-time. We instantiated RNNs by taking into account only the binary or weighted topology of BNNs. The results indicate that no differences are observed when using binary or weighted topologies. Evidently, the BNN to RNN extrapolation strategies and RNN architectures examined here are by no means comprehensive and, consequently, approaches that incorporate further biologically realistic features are needed, for instance, by incorporating different classes of excitatory and inhibitory neurons into RNNs.

### 6.3. Bio-instantiated RNNs from human and non-human animal brain networks

We used empirical data describing the network of the brain of diverse animals, that is, humans and monkeys. This comparative *in silico* examination is important for three reasons. First, it allows us to assess the importance of network topology of neural systems found in nature, without a bias that would be entailed by an exclusive use of data from one species, e.g., humans. Thus, we were able to examine the impact of biological network topology in RNNs from a universal, cross-species standpoint. Second, experimental methods for mapping neural systems have a different degree of reliability. Thus, converging evidence from data from human and non-human animals, collected with different methods (Markov et al., 2012; Betzel & Bassett, 2018; Majka et al., 2020), highlight the robustness of our results. Third, the comparative approach situates the human brain, and the brain of other animals on the same plane. Thus, our method constitutes the basis of future *in silico* examinations of species-specific features that may bestow each animal with unique functional and behavioral capacities.

### 6.4. Limitations and future directions

Our results demonstrate what strategies for creating bio-instantiated RNNs lead to RNNs with biological network topology without sacrificing performance. However, such network topology, *in the current setup and mode of evaluation*, does not convey any functional benefits, that is, achieving lower loss or converging faster to the minimum loss. Clearly, the RNNs as currently trained, possess several non-biologically plausible aspects, for instance, with respect to the training algorithm (backpropagation-through-time) and activity functions (ReLU, tanh). While in the current experiments we focused on ways to bestow RNNs with biologically realistic network topology, future studies should go beyond rate models and examine such network topology in a model with spiking neurons and appropriate, more biologically realistic training algorithms (Tavanaei et al., 2019). We should note that the effect of network topology, as shown in the context of Echo State Networks with threshold neurons (Rodriguez et al., 2019), may be enhanced in such context. In addition, biologically motivated principles, such as Dale’s principle, could be incorporated into the RNNs to enhance their realism, an endeavor rendered possible by recent methodological contributions (Song et al., 2016; Cornford et al., 2021). Lastly, the current mode of evaluation of bio-instantiated RNNs tests performance on isolated tasks. Extending the way that performance of bio-instantiated RNNs is evaluated could entail the incorporation of the wiring cost of the system (Suarez et al., 2020), or the benefit of bio-instantiated RNNs in e.g., transfer learning. Overall, the illustrated approach opens new ways to examine the intersection of ANNs and BNNs at the network topology level, with the capacity to enrich artificial systems with empirically discerned neurobiological properties. Such endeavor may create magnificent hybrid beasts, as well as some Chimeras.

## Acknowledgements

Funding is gratefully acknowledged: AG: Deutsche Forschungsgemeinschaft (SPP 2041 2888/2-2) FD: Deutscher Akademischer Austausch Dienst (DAAD), CCH: Deutsche Forschungsgemeinschaft (SFB 936/A1; TRR 169/A2; SPP 2041/HI 1286/7-1, HI 1286/6-1), the Human Brain Project (SGA2).

